# CD8^+^ T cell response promotes viral clearance and reduces chances of severe testicular damage in mouse models of long-term Zika virus infection of the testes

**DOI:** 10.1101/2024.01.22.575592

**Authors:** Rafael K. Campos, Yuejin Liang, Sasha R. Azar, Judy Ly, Vidyleison Neves Camargos, E. Eldridge Hager-Soto, Eduardo Eyzaguirre, Jiaren Sun, Shannan L. Rossi

## Abstract

Zika virus (ZIKV) causes human testicular inflammation and alterations in sperm parameters and causes testicular damage in mouse models. The involvement of individual immune cells in testicular damage is not fully understood. We detected virus in the testes of the interferon (IFN) α/β receptor^-/-^ A129 mice three weeks post-infection and found elevated chemokines in the testes, suggesting chronic inflammation and long-term infection play a role in testicular damage. In the testes, myeloid cells and CD4^+^ T cells were absent at 7 dpi but were present at 23 days post-infection (dpi), and CD8^+^ T cell infiltration started at 7 dpi. CD8^-/-^ mice with an antibody-depleted IFN response had a significant reduction in spermatogenesis, indicating that CD8^+^ T cells are essential to prevent testicular damage during long-term ZIKV infections. Our findings on the dynamics of testicular immune cells and importance of CD8^+^ T cells functions as a framework to understand mechanisms underlying observed inflammation and sperm alterations in humans.

## 1 INTRODUCTION

Zika virus (ZIKV) is a positive-strand RNA virus of the *Flaviviridae* family that was discovered in Uganda in 1947 and is transmitted primarily by mosquito bites. However, Foy and colleagues proposed an additional mode of transmission, sexual transmission, when a likely case was identified in the United States in 2008-2009, involving a man that had returned from Senegal in 2008-2009^1–2^. Sexual transmission was confirmed during the 2015-2017 ZIKV epidemics^3^, which together with animal experiments^4^, confirmed that ZIKV could be sexually transmitted^1, 3, 5–7^. Confirmation of this mode of transmission raised numerous concerns with human health implications, including effects it could have on ZIKV circulation during epidemics, enhanced concerns of ZIKV congenital syndrome, as well as testicular damage. ZIKV infectious particles or ZIKV RNA were present in semen and vaginal secretions long after the initial infection^8–9^. In semen, ZIKV infectious particles were detected for up to 69 days^10^, and ZIKV RNA has been detected for up to 370 days after initial infection^11^. Men that shed ZIKV RNA for long periods have signs of male reproductive tract inflammation, with higher leukocyte counts^12^, elevated cytokine levels^12^, and several alterations in sperm^13^. Since ZIKV does not commonly cause fatal disease, acquiring tissue samples of people recently infected with ZIKV is challenging.

Using animal and cell culture models, components of the male reproductive tract shown to be a site of long-term replication and inflammation include the epididymides^14–16^, the prostate^17–18^, and the testes ^19–21^, leading to reduced fertility in mouse models ^21^. Studies using ZIKV-infected NHP^22^ and mouse models ^20–21^ have also showed ZIKV infection can lead to histopathologic lesions in the testes, which is a health concern for infected men in addition to impacting fertility. The testes are the site of spermatogenesis and may play a key role in ZIKV-caused infertility as well as viral maintenance for sexual transmission. Studies have shown that ZIKV can establish long-term infections in Sertoli cells, germ cells^23^, and spermatogonia^23^.

The testes are an immune-privileged organ with a delicate balance maintained by the blood-testis barrier (BTB) and regulatory, tolerogenic, immune cells, including macrophages, dendritic cells, mast cells, cluster of differentiation (CD)4^+^ and CD8^+^ T cells ^24–25^. Macrophages are the major population of cells in the testes, which represents approximately 20% of cells in the interstitial space under physiological conditions. Macrophages and dendritic cells have been found to be important targets of ZIKV visceral replication^26^. In addition to immune cells native to the testes, cell infiltration and inflammation have also been shown to be triggered by ZIKV infection in animal models^20, 27^, and one cell population confirmed to invade the testes is CD8^+^ T cells^28^. Macrophages and dendritic cells were implicated as important initial targets of replication which may help the virus invade the testes ^26, 29^. However, the dynamics of infiltration of immune cells in the testes and their roles in clearing ZIKV infection are not well understood.

Using mouse models, we have investigated cytokine levels and immune cells involved in the response against ZIKV long-term testicular infection and the role of CD8^+^ T cells in clearing ZIKV from testicular tissue and reducing the odds of testicular damage.

## 2 RESULTS

### 2.1 ZIKV long-term infection of the testes in human primary Sertoli cells and mouse models

Sertoli cells comprise the BTB and are essential for maintaining seminiferous tubule (ST) architecture. Both murine 15P-1 and primary human (hp) Sertoli cells support ZIKV PRVABC59 infection (Figure 1A). hpSertoli cells support higher viral titers in the supernatant (6 log_10_ plaque-forming units (PFU)/ml). In both of these cell types, viral titers are maintained for at least 5 days while no substantial cell death is observed. For hpSertoli cells, we measured that infection at a multiplicity of infection (MOI) of 0.1 for 5 days does not significantly lower cell metabolic activity (Figure 1B). We then performed *in vivo* experiments using the A129 mouse model, infecting these mice with a target dose of 3 log_10_ PFU of ZIKV intraperitoneally. Viremia at 2 dpi confirmed that all animals were infected (Figure 1C). The weight of the testes was significantly reduced compared to that of the PBS-injected mice (Figure 1D). Several testes from infected mice had apparent testicular damage with diffuse tubular necrosis upon histological examinations (Figure 1E). This appears to contrast with the resistance to cell death upon infection observed in hpSertoli cells in culture (Figure 1A-B). It is possible that the immune system contributes to the testicular damage by disrupting the testicular microenvironment.

**Figure 1.**
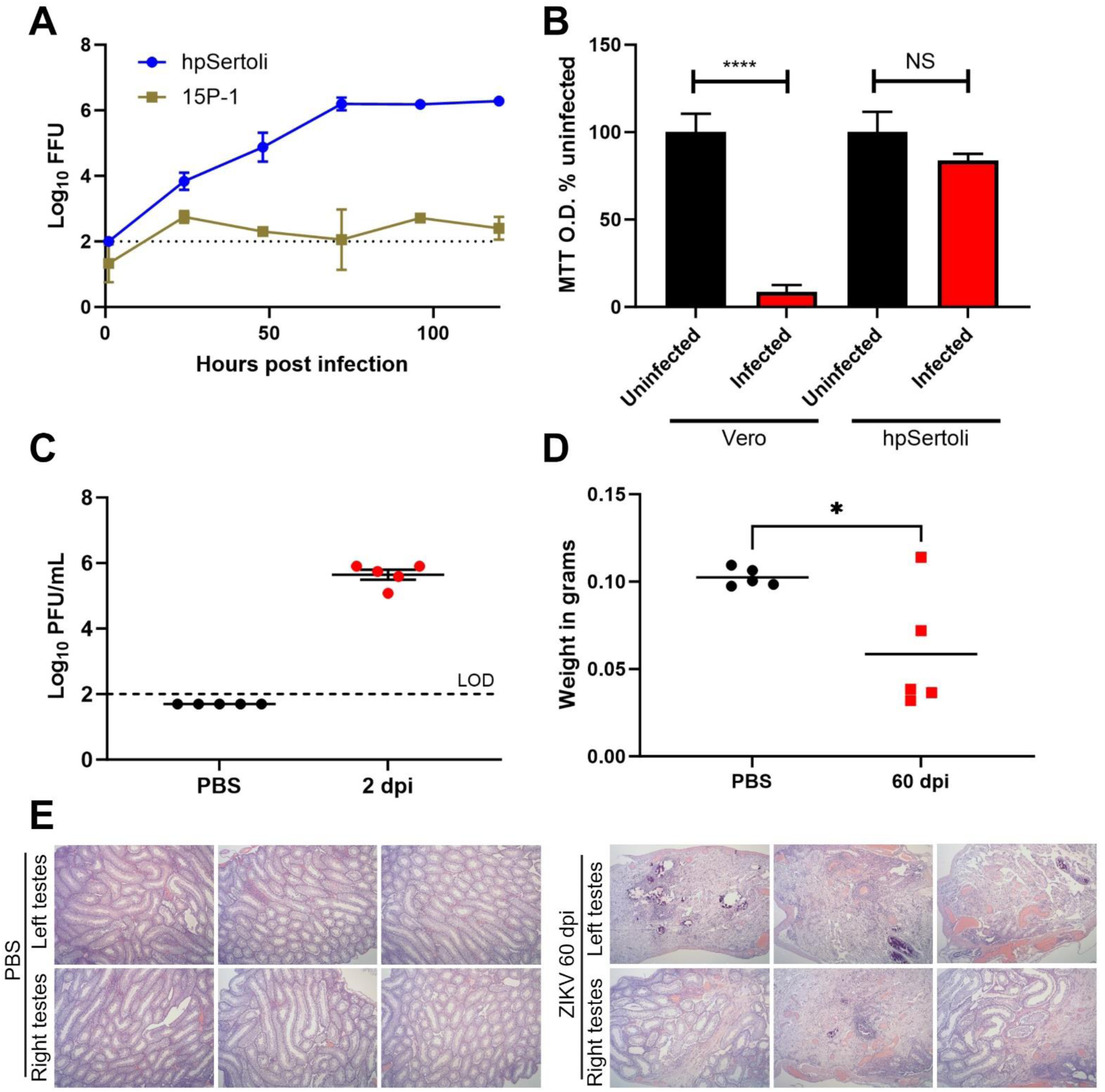
Primary human Sertoli cells support long-term infection of ZIKV with minimal cell death, but widespread damage is observed in testes of A129 mice. A. ZIKV PRVABC59 growth curve on primary human Sertoli (hpSertoli) cells or mouse Sertoli (15P-1) cells at an MOI of 0.1, the supernatant was collected at each time point and titrated B. Sertoli cells show minimal cell death when MTT assay is used as a proxy for cell viability. For A. and B., bars show mean and error bars represent standard deviation. Statistical significance was assessed with one-way ANOVA Sidak’s multiple comparison correction. C. Viremia in A129 mice at 2 dpi. D. A129 mice testicular weight 60 dpi with ZIKV. Statistical significance was assessed using a two-tailed t-test. Lines represent the mean. E. Histopathological damage observed in mice testes 60 dpi with ZIKV. *, p < 0.05; ****, p < 0.0001.

### 2.2 ZIKV long-term infection of the testes in the A129 mouse model induces local cytokine and chemokine expression

To investigate whether long-term ZIKV infection of the testes induced local immune system activity, we infected A129 mice with a target dose of 3 log_10_ PFU of ZIKV. We observed that the mice showed acute signs of disease, such as lethargy, ruffled fur, and had significant body weight loss, with peak weight loss at 10 days post-infection (dpi) before recovery (Figure 2A). All mice developed viremia at 2 dpi with an average around 6 log_10_ PFU/ml, whereas none had detectable viremia three weeks after infection (Figure 2B), suggesting systemic infection had been cleared. In contrast, at 21 dpi we detected viable virus in the testes of 3 out of 6 mice, with 2 of these mice having virus in both of the testes, and the highest titer detected being 7 log_10_ PFU/mg of tissue (Figure 2C). Cytokines and chemokines presented in testicular macerates and plasma at this time were analyzed on a multiplex bioassay (BioPlex). We detected a significant but moderate (< 5-fold) increase in interleukin (IL)-3 and G-CSF in the plasma (Figure 3A). In testicular homogenates, we detected a significant and drastic change in the pro-inflammatory cytokines IL-1α and IL-12(p40), and in the chemokines C-C motif chemokine ligand (CCL)-3, CCL-4 and CCL-5 (Figure 3B). The elevated levels of chemokines suggest that immune cells are being recruited in the testes, and chronic inflammation may occur three weeks after the initial infection.

**Figure 2.**
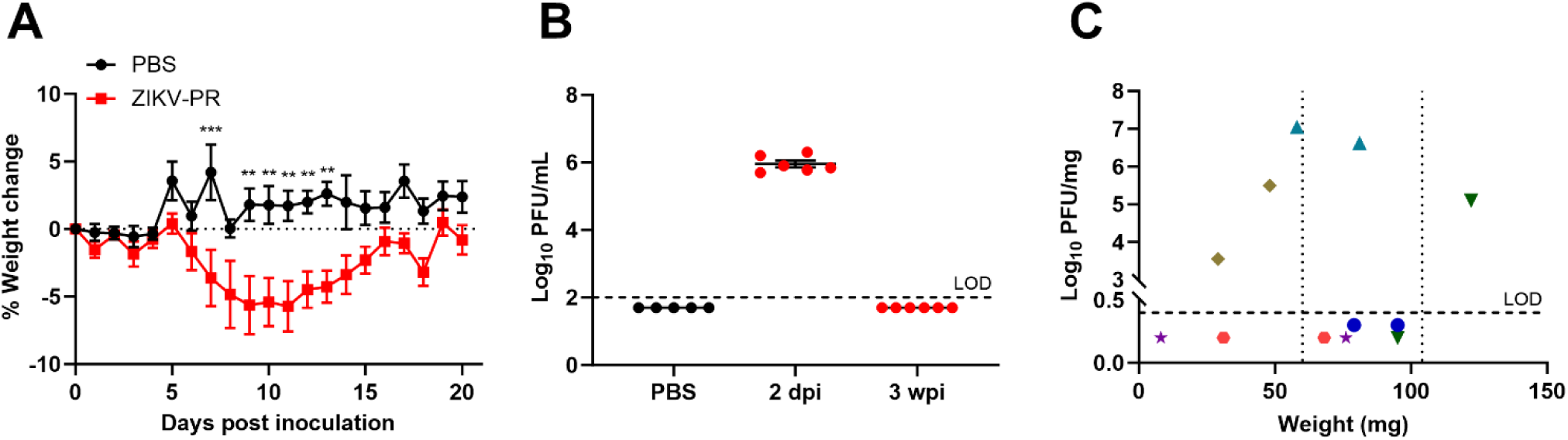
ZIKV replication in testes of A129 mice after clearance of systemic infection. A. Weights of A129 mice infected with ZIKV PRVABC59. Statistical significance was assessed with two-way ANOVA with Tukey’s multiple comparison correction. Error bars represent standard error of the mean. B. Virus detection in the plasma of A129 mice, uninfected, or at 2 or 23 dpi. Lines represent the mean. C. Testicular weight of the mice (x-axis) and viral titer in the testes (y-axis) at 23 dpi. Same colors indicate that the testes are from the same mouse. Bars show mean and error bars represent standard error of the mean. Vertical dotted lines indicate the range of the testicular weight of PBS-injected mice. Horizontal dotted lines represent the limit of detection. ***, p < 0.001.

**Figure 3.**
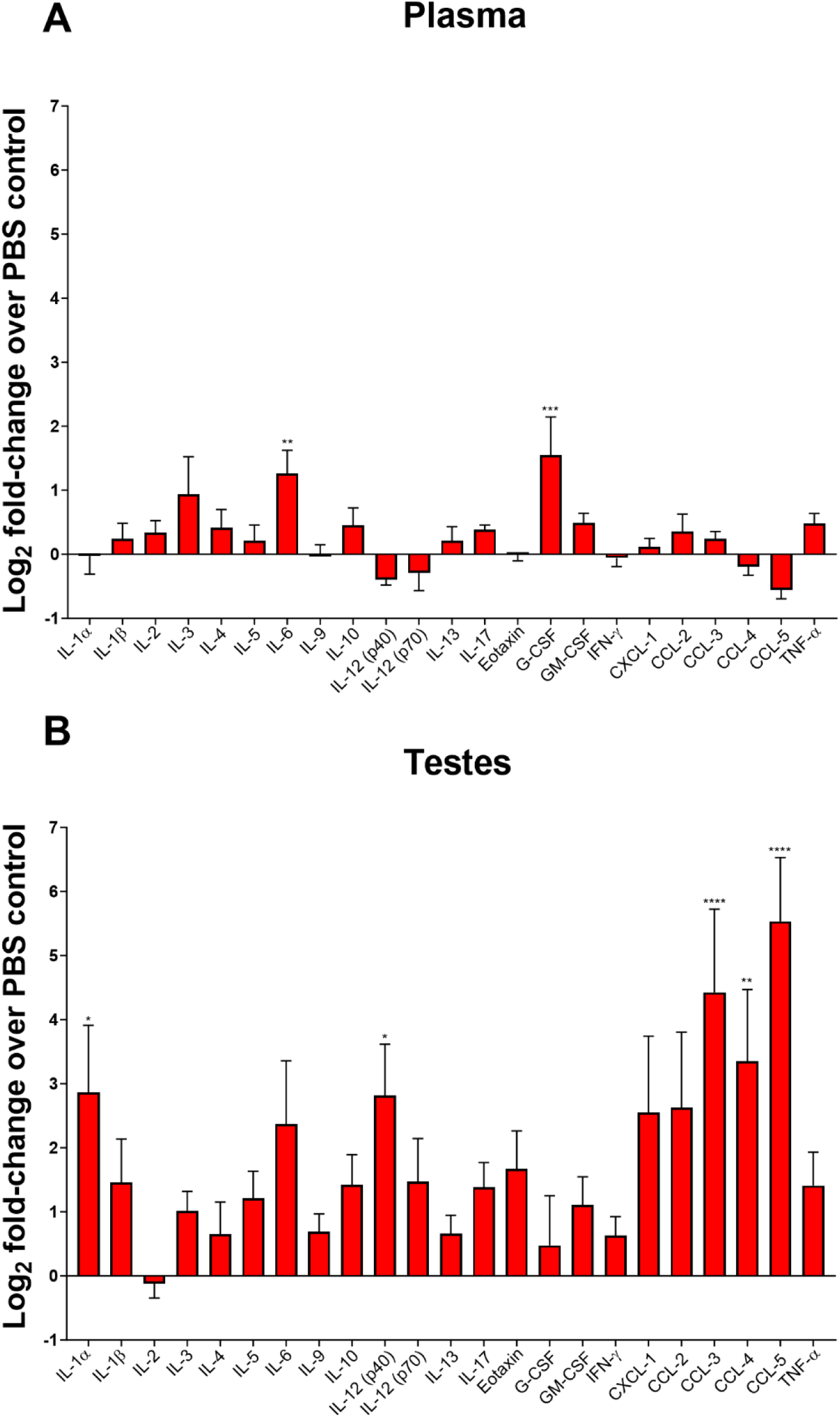
Long-term ZIKV infection of A129 mice induces local activation of cytokines and chemokines. A. Log_2_ of the fold-change of cytokines and chemokines measured in the plasma of ZIKV-infected mice. B. Fold-change induction of cytokines and chemokines in the testes of ZIKV-infected mice. Statistical significance was assessed with one-way ANOVA Sidak’s multiple comparison correction in between PBS control groups (N = 5) and the infected groups (N = 6), for each cytokine and chemokine. Bars show mean and error bars represent standard error of the mean. *, p < 0.05; ****, p < 0.0001.

### 2.3 Myeloid cells are present in the lymphoid organs primarily at 7 dpi and in the testes at 23 dpi

To investigate immune populations in the context of long-term ZIKV infection of the testes, we infected A129 mice with a target dose of 3 log_10_ PFU of ZIKV intraperitoneally. We observed similar weight loss, disease, and recovery of visible disease signs as in the previous experiment. In addition to collecting testicular tissue, to assess the systemic presence of immune cells in the different lymphoid organs, we collected the spleen, the mesenteric lymph nodes, and the inguinal lymph nodes. Samples were processed for flow cytometry, and data were collected on immune cell populations at 7 and 23 dpi. We counted the total number of cells in the testes and the lymphoid organs and found that the spleen and the mesenteric lymph nodes had increased total cell counts at 7 dpi. Cells from the lymph nodes were significantly reduced from 7 dpi to 23 dpi, but total cells in the spleen remained elevated at the 23 dpi timepoint (Figure 4A-C). In contrast to the lymphoid organs, the total number of cells in the testes was not significantly different (Figure 4D). In the spleen, the mesenteric and inguinal lymph nodes, there was an increase in macrophages and dendritic cells at 7 dpi, compared to the organs of PBS-inoculated controls (Figure 4A-C). Neutrophils were also elevated in the spleen and the inguinal lymph nodes at 7 dpi but did not reach statistical significance in the mesenteric lymph nodes (Figure 4A-C). At 23 dpi, the levels of these cells were either similar to 7 dpi or decreased in the lymphoid organs (Figure 4A-C). In contrast to the lymphoid organs, myeloid cells were not present in the testes at 7 dpi but were highly increased at 23 dpi (Figure 4D). The presence of myeloid cells in the testes late in infection indicates chronic inflammation.

**Figure 4:**
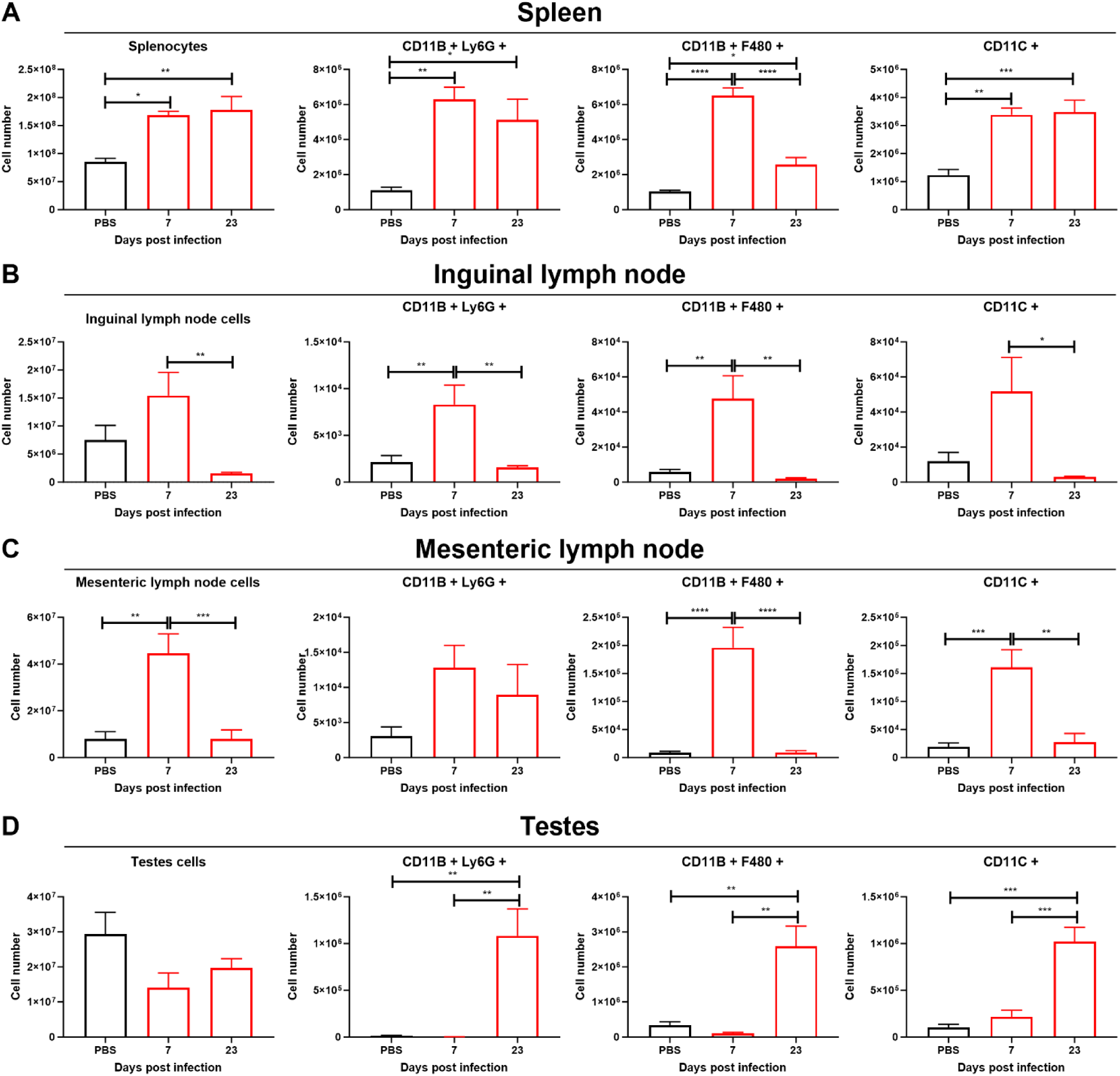
Macrophages, neutrophils and DCs infiltrate testes at 23 dpi. Cell count and myeloid cells for A. the spleen, B. the inguinal lymph nodes, C. the mesenteric lymph nodes and D. testes. Red bars show ZIKV-infected mice, black bars denote uninfected baseline controls. Statistical significance was assessed with one-way ANOVA Sidak’s multiple comparison correction. Bars show mean and error bars represent standard error of the mean. *, p < 0.05; **, p < 0.01; ***, p < 0.001; ****, p < 0.0001.

### 2.4 CD4^+^ T cells but not CD8^+^ T cells recruitment to the testes are delayed in comparison to the lymphoid organs

As T cells are an integral part of the immune response against ZIKV, we assessed CD4^+^ T cells at 7 and 23 dpi in these ZIKV-infected mice. We found that CD4^+^ effector T cells (CD44^hi^, CD62L^lo^) were increased in the lymphoid organs at 7 dpi, and then decreased back to original levels at 23 dpi in the inguinal and mesenteric lymph nodes but not in the spleen (Figure 5A-C). In contrast, CD4^+^ effector T cells were not significantly increased in the testes at 7 dpi but were increased at 23 dpi (Figure 5D). Effector T cells prominently express CXCR3, which plays a crucial role in regulating T cell trafficking and function ^30–31^. In the lymphoid organs, CD4^+^ CXCR3^+^ T cells were increased at 7 dpi, and decreased again in the lymph nodes at 23 dpi, but remained elevated in the spleen (Figure 5A-C). In the testes, CD4^+^ CXCR3^+^ T cells were not increased at 7 dpi, but were drastically increased at 23 dpi, similar to what was observed for myeloid cells. IFN-γ plays a chief role in virus immunity and countering ZIKV infections. We evaluated CD4^+^ IFN-γ^+^ T cells and found that it was broadly consistent with the presence of CD4^+^ effector T cells, being increased in the lymphoid organs but not in the testes at 7 dpi (Figure 5A-D). At 23 dpi, CD4^+^ IFN-γ^+^ T cells were decreased in the lymphoid organs in comparison to 7 dpi (Figure 5A-C), but were strongly increased in the testes, suggesting a chronic activation of the IFN antiviral response in testicular tissues. IL-17 is an important factor related to allergies and autoimmunity, and functions in many biological processes, including chronic inflammation. IL-17 is thought to be produced mainly by Th17 cells, a subset of CD4^+^ T cells. We found that CD4^+^ IL-17^+^ T cells were increased at 7 dpi in the spleen and the mesenteric lymph nodes but not in the testes and the inguinal lymph nodes, which drain from the testes (Figure 5 A-D). At 23 dpi, CD4^+^ IL-17^+^ T cells were no longer statistically increased compared to uninfected controls, and the inguinal lymph nodes continued without any increase in these cells. In contrast, the testes had a large population of CD4^+^ IL-17^+^ T cells present at this time point. We also observed a population of possible memory cells CD4^+^ CD44^hi^ CD62L^hi^ which increased in the lymphoid organs at 7 dpi and in the testes at 23 dpi and could have implications for a secondary testicular infection of ZIKV (Figure S2).

**Figure 5:**
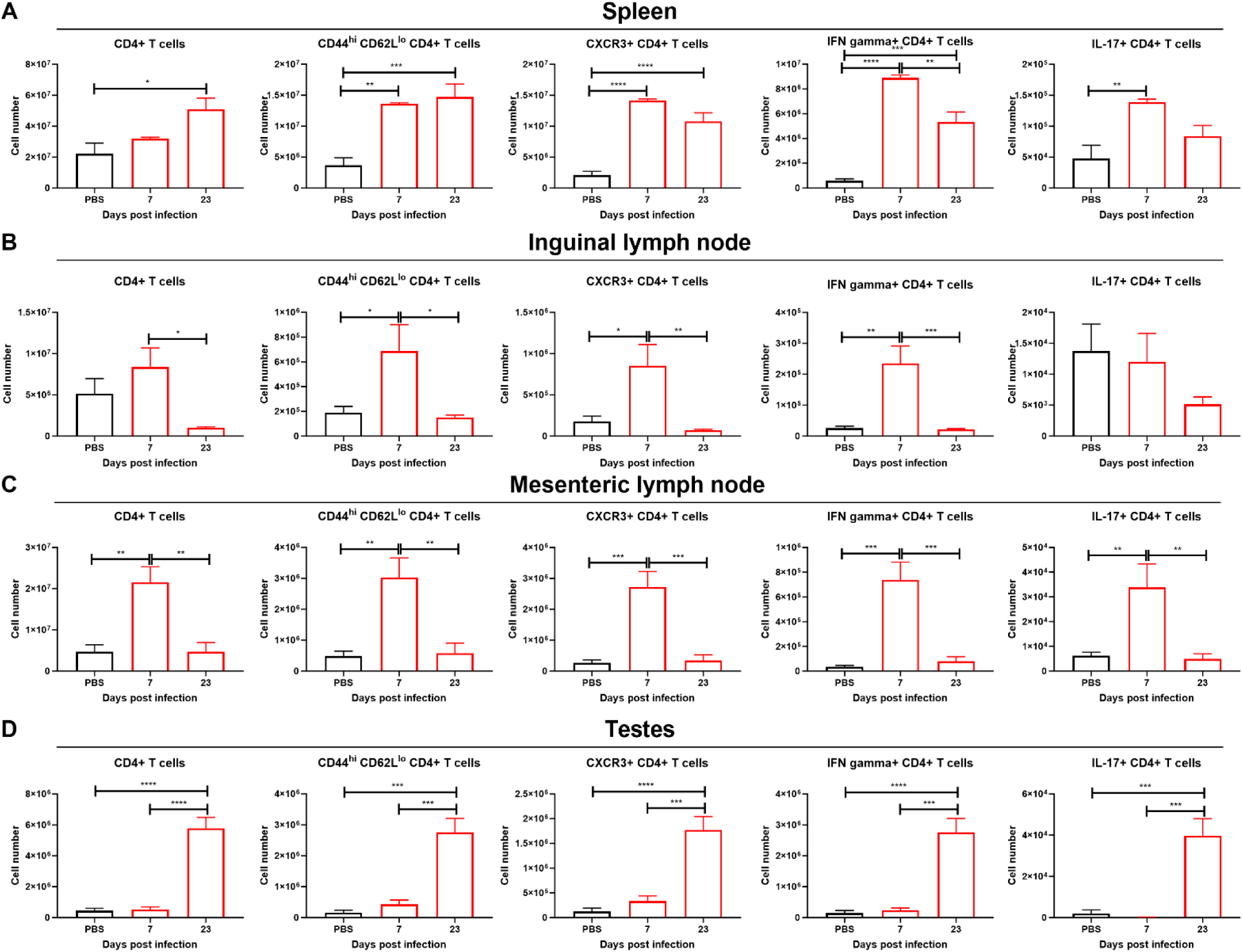
CD4^+^ T cells accumulate in testes at 23 dpi. CD4^+^ T cells for A. the spleen, B. the inguinal lymph nodes, C. the mesenteric lymph nodes and D. testes. Red bars show ZIKV-infected mice, black bars denote uninfected baseline controls. Statistical significance was assessed with one-way ANOVA Sidak’s multiple comparison correction. Bars show mean and error bars represent standard error of the mean. *, p < 0.05; **, p < 0.01; ***, p < 0.001; ****, p < 0.0001.

In all organs tested, effector CD8^+^ T cells (CD44^hi^, CD62L^lo^) were increased at 7 dpi, earlier than other cell types tested (Figure 6A-D). At 23 dpi, effector CD8^+^ T cells and CD8^+^ CXCR3^+^ T cells were decreased compared to 7 dpi in the lymphoid organs and were no longer significantly higher than uninfected in the testes (Figure 6A-D). That was also the case for CD8^+^ IFN- γ^+^ T cells in the lymphoid tissues but not for the testes (Figure 6A-D). In testicular tissue, CD8^+^ IFN-γ^+^ T cells were not significantly increased at 7 dpi, but were significantly higher at 23 dpi (Figure 6D). This indicates that at 7 dpi in the testes, there are many effector CD8^+^ T cells, but these are not good producers of IFN- γ^+^, and only later in long-term ZIKV infection there is presence of CD8^+^ IFN-γ^+^ T cells. For both CD4^+^ and CD8^+^ T cells, we found that programmed Death-1 (PD-1), an inhibitory receptor induced in activated T cells was consistent with the measurements of effector cells using CD44 and CD62L (Figure S3).

**Figure 6:**
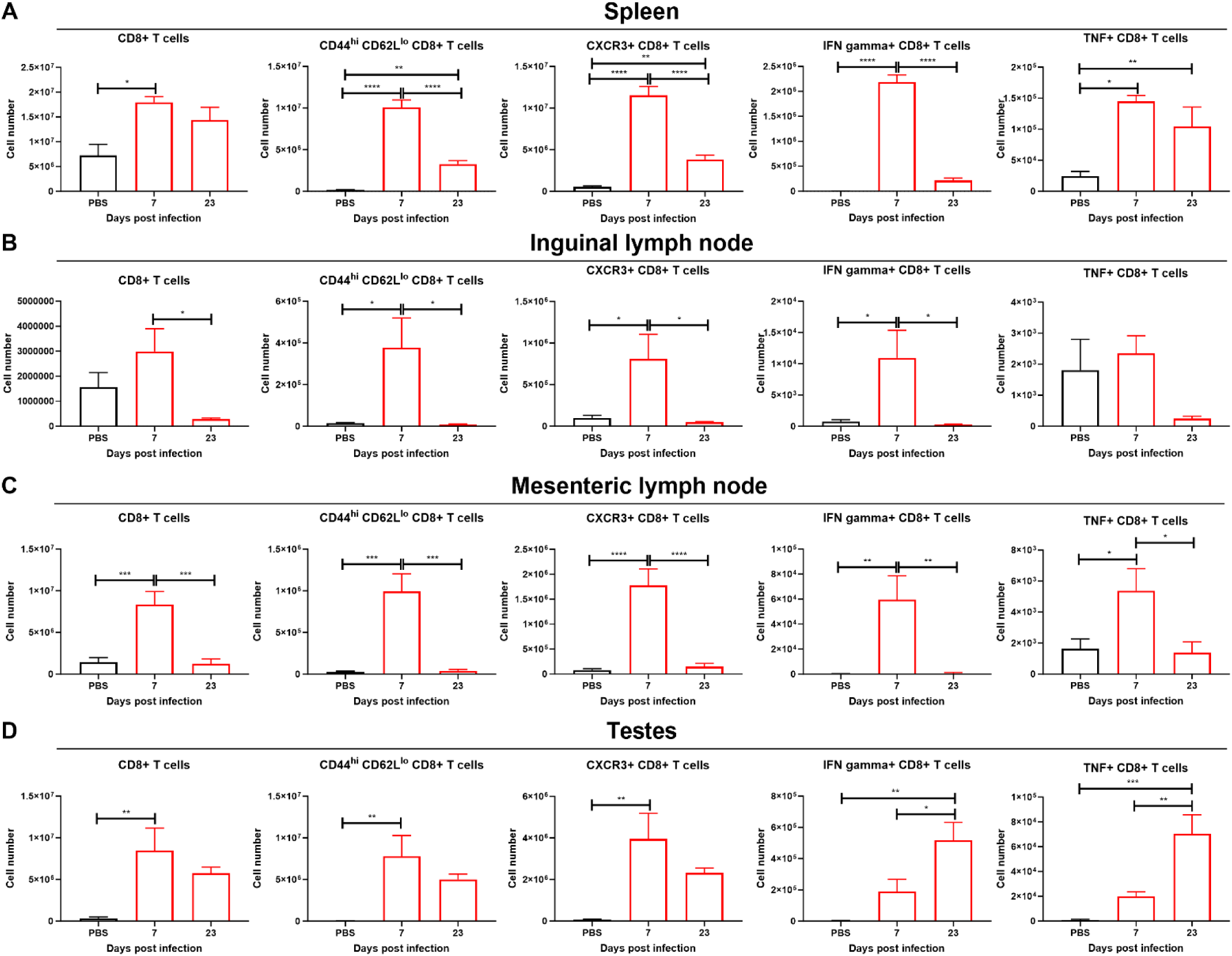
CD8^+^ T cells accumulate in testes at 7 and 23 dpi and in lymph nodes at 7 dpi. CD8^+^ T cells for A. the spleen, B. the inguinal lymph nodes, C. the mesenteric lymph nodes, D. testes. Red bars show ZIKV-infected mice, black bars denote uninfected baseline controls. Statistical significance was assessed with one-way ANOVA Sidak’s multiple comparison correction. Bars show mean and error bars represent standard error of the mean. *, p < 0.05; **, p < 0.01; ***, p < 0.001; ****, p < 0.0001.

### 2.5 C57BL/6J CD8^-/-^ have decreased spermatogenesis compared to wild type control mice

As the flow cytometry data indicated a critical role for T cells, we next sought to assess their role in the establishment of viral infection or viral testicular clearance, and if they influenced the severity of testicular damage in long-term ZIKV infection. Two mouse models, the A129 mice (IFNα/βR^-/-^) or the immunocompetent C57BL/6J mice, were used; cohorts of mice were treated with anti-mouse CD3ε F(ab’)2 fragments or control f(ab’)2 fragments of polyclonal hamster IgG (5 injections of 100 µg each of anti-CD3 or control treatments on the following dpi: -3, -1, 2, 7, and 14). Because ZIKV cannot efficiently infect C57BL/6J mice, these mice were additionally transiently injected with 1.5 mg/mouse of IFNAR-blockading antibody one day before infection to allow ZIKV to infect the mice tissues and reach the testes. Mice were infected with ZIKV as previously done. To confirm the depletion of CD3^+^ cells, we measured CD3 in the blood by flow cytometry on 2 and 18 dpi. In A129 mice, the depletion was approximately 70% at 2 dpi, and >99% at 18 dpi, and in C57BL/6J, the depletion was approximately 80% in both timepoints (Figure S4A-B). However, in both mouse models, there was no significant difference for percentage of ST with spermatogenesis or testicular weights between infected control and CD3 depleted mice (Figure S4C-H).

As CD8^+^ T cells were the only cell type detected abundantly in the testes at 7dpi, we hypothesized that these cells might contribute to the early immune reaction observed, leading to the recruitment of other immune cells. To test that hypothesis, we intraperitoneally infected wild type (WT) C57BL/6J mice or its CD8^-/-^ counterpart with a 5 log_10_ PFU/mouse target dose. In this experiment, to obtain a more permissive testicular damage phenotype, we injected the mice twice with 1.5 mg/mouse of IFNAR-blockading antibody one day before infection and five dpi. To quantify testicular damage in the hematoxylin-eosin-stained histology slides (Figure 7A), a board-certified pathologist blinded to the samples quantified the percentage of the ST with spermatogenesis in each slide. The CD8^-/-^ mice had significantly decreased percentage of ST with spermatogenesis compared to the WT C57BL/6J mice (Figure 7A-C), suggesting the presence of CD8^+^ T cells is key to avoiding or reducing the severity of the testicular damage. We evaluated the slides by immunofluorescence (Figure 7D) and detected ZIKV envelope protein in 0 out of 10 WT mice and 6 out of 10 CD8^-/-^ at 23 dpi (Fisher’s exact test, p =0.01). We observed presence of viral envelope protein in cells positive for DDX-4 (marker of spermatogonia, spermatocytes, and round spermatids) and in cells negative for DDX-4, suggesting ZIKV infects various cell types in the ST (Figure 7E).

**Figure 7:**
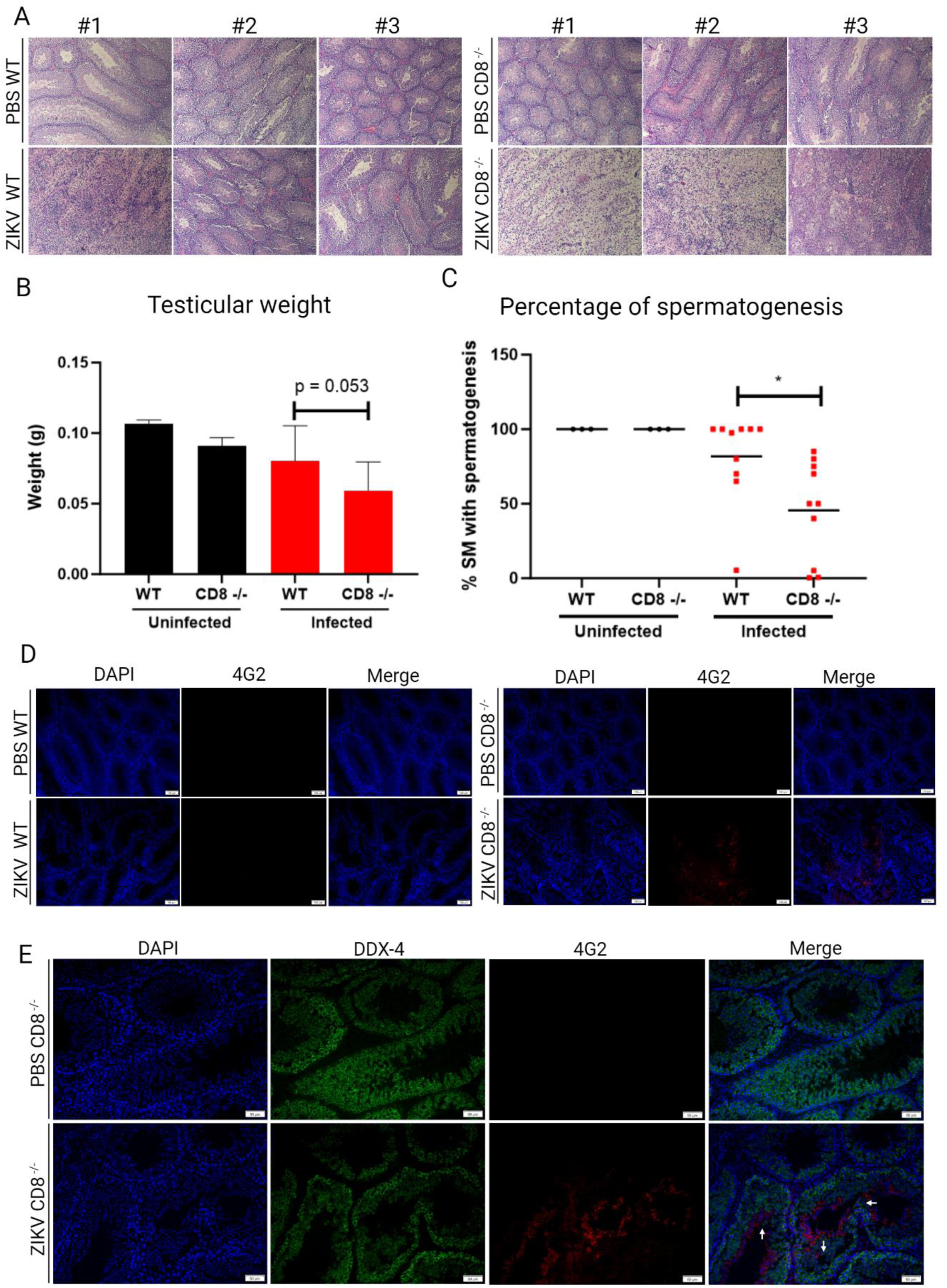
CD8^+^ T cells are important to prevent significant testicular damage upon long-term ZIKV infection in C57BL/6J mice transiently treated with anti-IFN antibodies. A. Histology of WT C57BL/6J mice or CD8^-/-^ mice testes. B. Testicular weight. Statistical significance was assessed with one-way ANOVA Sidak’s multiple comparison correction in between WT and CD8^-/-^ conditions, either uninfected or infected. Bars show mean and error bars represent standard deviation. C. Percentage of spermatogenesis quantified from the histology by a pathologist blinded to the samples. Statistical significance was assessed using a two-tailed t-test. Lines represent the mean. D. Immunofluorescence of testicular samples showing cell nuclei (blue), ZIKV envelope protein (red), and DDX-4 (green). The white arrows show cell detachment from wall of the ST. *, p < 0.05.

## 3 DISCUSSION

ZIKV has caused large epidemics worldwide, but most notably across the Americas in 2015 and 2016. This epidemic had an enormous medical, societal, and economic cost, and was associated with previously poorly characterized disease manifestations including congenital Zika syndrome and Guillain Barre syndrome. The repercussions of this outbreak are still felt today, with communities and healthcare systems continuing to bear the burden of its long-term consequences. During that epidemic, researchers confirmed earlier reports ^1, 7^ that ZIKV could be sexually transmitted, as well as cause chronic inflammation in the male reproductive tract ^12^. Animal model experiments have implicated the testes as a site of prolonged infection and inflammation ^19–20^. Although in vasectomized men, detection of ZIKV is still possible, suggesting ZIKV may infect tissues distal from the vas deferens, such as the prostate, seminal vesicles, and bulbourethral glands^32^. The testes remain as a key site of viral replication and long-term maintenance, and impact on this organ may affect fertility.

For other viral infections of the testes, such as mumps virus, cytokines play a role disrupting the BTB for the virus to first enter the testicular environment^33^. At three weeks post-infection, we found that IL-6 and G-CSF cytokines were upregulated in the plasma (Figure 3A). Higher IL-6 has been associated with severity of COVID-19 in humans^34^. It is possible that earlier in infection, more cytokines were upregulated in the plasma, leading to a breakdown of the BTB that may accelerate ZIKV entrance in the testicular environment. Cytokines known to disrupt the BTB integrity include IL-6^35^, which was detected in plasma of the A129 mice three weeks after initial infection (Figure 3A), TNF^36^, IL-1^37^, TGF-β3^36^. Some cytokines are also able to cross the BTB, such as G-CSF^38^, which was also detected (Figure 3A). However, since ZIKV can directly infect Sertoli cells (Figure 1), it may be able to partly bypass the BTB. In the testes, we found that high levels of chemokines CCL-3, CCL-4 and CCL-5 were present, suggesting recruitment of immune cells and chronic inflammation. Several cytokines were also increased over 10-fold in individual infected mice in comparison to uninfected controls, but when data from multiple animals was pooled for statistical analysis, it did not reach significance.

In a ZIKV infection model using A129 mice we only detected CD8^+^ T cells at the earlier timepoint of 7 dpi. However, it is possible that myeloid cells had already infiltrated the tissue at an even earlier timepoint but were no longer present at 7 dpi. Some myeloid cells, such as macrophages and dendritic cells have been shown to be particularly important in early infection for ZIKV to establish testicular infection^26, 29^. We found presence of myeloid cells, CD4^+^ T cells and CD8^+^ T cells in the testes at 23 dpi. This contrasted with the lymphoid organs, which had an increase in most of the immune cells at 7 dpi, but the levels of these cells then decreased or stayed similar at 23 dpi, possibly reflecting the systemic viral clearance at that timepoint. We treated A129 or C57BL/6J mice (transiently depleted for IFN signaling using antibodies) with anti-CD3, with the purpose to deplete CD4^+^ and CD8^+^ T cells. We did not observe statistically significant differences between control and CD3-depleted mice in testicular weight, histological assessments, or viral load in the testes. This was in strong contrast to the experiment assessing C57BL6/J CD8^-/-^ mice, which completely lack CD8^+^ T cells and had increased testicular damage. The discrepancy may arise from anti-CD3 treatments depleting not only CD8^+^ T cells but also CD4^+^ T cells and other cell types^39^. It is possible these cells contribute to exacerbated immune response leading to more testicular damage, and that, conversely, some reduction in their population improves the immune response profiles and mitigates testicular damage, counteracting the effects of CD8^+^ T cell depletion. Alternative explanations for these differences may be that some presence of CD8^+^ T cells is required to avoid severe testicular damage, but that having reduced levels of CD8^+^ T cells may not present as a particular issue, or due to differences in the regimen of anti-IFN antibodies in between these two experiments. Studies to separate these possibilities and investigate the role of other cells of the immune system seem warranted.

In agreement with previous reports that CD8^+^ T cells are protective in the context of systemic infections^40^, we show these cells are also important in the context of testicular infection, for both viral clearance and reduced chances of testicular damage. A similar protective role for CD8^+^ T cells has been shown in the context of the central nervous system, which like the testes is immune privileged^31^. CD8^+^ T cells recruited in a CXCR3-dependent manner can effectively control the virus in the brain if they arrive early after viral invasion in sufficient numbers and appropriate differentiation state ^31^.

In summary, our results offer insights into the kinetics of immune cell infiltration in the testes and the role of CD8^+^ T cells in testicular damage in the context of long-term ZIKV testicular infections. We observed early infiltration of CD8^+^ T cells in the testes, and that presence of CD8^+^ T cells promote viral clearance and reduce chances of testicular damage.

## 4 METHODS

### 4.1 Cells and viruses

Vero CCL-81 and 15-P1 cells (American Type Culture Collection) were grown in Dulbecco’s minimal essential media (DMEM, Gibco, Thermo Fisher Scientific) with 10% fetal bovine serum, 200 U/mL penicillin and 200 mg/mL streptomycin (DMEM, Gibco, Thermo Fisher Scientific). Primary human Sertoli cells were purchased from and grown in Sertoli cell growth media (IX Cells Biotechnologies) supplemented with 10% fetal bovine serum (IX Cells Biotechnologies), Sertoli cell growth supplement according to the manufacturer’s recommendations (IX Cells Biotechnologies), and antibiotic-antimycotic according to the manufacturer’s recommendations (IX Cells Biotechnologies). Sertoli cells were used in passage 3 after acquisition. ZIKV PRVABC59 was received from the World Reference Center for Emerging Viruses and Arboviruses, UTMB (WRCEVA) at Vero passage 4. The virus underwent two additional Vero cell passages to generate the stocks utilized in these studies, as previously described ^41^.

### 4.2 Virus titration by plaque assays

Plaque assays were performed on Vero monolayers on 12-well plates as previously described. Briefly, samples were diluted serially and used to infect Vero monolayers and after 1 h rocking at 37°C, overlayed with DMEM containing 0.8% methylcellulose. Plates were incubated at 37 °C in an atmosphere with 5% CO2 for approximately 108 h before fixation with a 10% formaldehyde solution. The cells were fixed with a solution of 0.2% crystal violet in 30% methanol to visualize the plaques. Data are shown as PFU/ml or PFU/mg, with the limit of detection (LOD) indicated on each graph by a dotted line. For PFU/mg, each sample had slightly different LODs, and the dotted line on the graph represents the highest LOD. Values below the LOD were set to half of the LOD for statistical and graphing purposes.

### 4.3 Animal infections

A129 mice are maintained in sterilized caging in a breeding colony at UTMB. C57BL/6J and CD8^-/-^ (B6.129S2-Cd8atm1Mak/J) were obtained from Jackson Laboratory. Animals were ear-notch identified. All animal manipulations were done following an approved Institutional Animal Care and Use Committee (IACUC). The animal weights were taken up to 14 dpi daily and every third day thereafter. 9–10-week-old A129 mice were infected intraperitoneally with a target dose of 3 log_10_ PFU/mouse. 8-9-week-old C57BL/6J and CD8^-/-^ were infected with a target dose of 5 log_10_ PFU/mouse. C57BL/6J and their CD8^-/-^ counterparts were treated with 1.5 mg/mouse IFNAR-blockading antibody (MAR1-5A3, Leinco) once (one day before infection, in studies shown in Figure S4) or twice (one day before infection and 5 dpi, in the study shown in figure 7). In the experiment shown in Figure 1, 5 mice were used as PBS controls, and 5 mice were used in the infection experimental condition. In the experiment shown in Figures 2 and 3, 5 A129 mice were used as PBS controls and 6 mice were used for the infected condition. In the experiment shown in Figures 4, 5, and 6, 3 mice were used in the PBS condition and 5 in the ZIKV-infected condition. In the experiments shown in Figure 7, 5 C57BL/6J mice were used in each condition, and 10 A129 were used in each condition. In the experiment shown in Figure 8, 3 WT and 3 CD8^-/-^ were used in the PBS conditions; 10 WT and 10 CD8^-/-^ were used in the infected groups. In the experiments shown in Figure S4, A129 or C57BL/6J mice were used. Two experiments were done with A129 mice, and the data combined, and one with C57BL/6J, each experiment using 5 mice per group. For experiment shown in Figure S4, 0.1 mg/mouse of anti-mouse CD3ε F(ab’)2 fragments (clone 145-2C11, BioXCell) or control f(ab’)2 fragments of polyclonal hamster IgG (BioXCell) were injected intraperitoneally on days -3, -1, 2, 7, and 14 relative to infection at day 0.

### 4.4 Flow cytometry and multiplex cytokine/chemokine assay

To measure levels of cytokines and chemokines in the plasma and the testes macerates of mice, the samples were prepared as instructed by the manufacturer and read using Bio-Plex Pro Mouse Cytokine 23-Plex (Bio-Rad, Hercules). Samples were run in technical duplicates, and the average was used. To quantify testicular cell populations, individual testes were macerated through 70-µm cell strainers and digested with 0.05% collagenase type IV (Thermo Fisher Scientific) in RPMI 1640 medium for 30 mins at 37°C. These samples were again passed through 70-µm cell strainers to prepare single-cell suspension. Samples of inguinal lymph nodes, mesenteric lymph nodes, and the spleen were passed through cell strainers in RPMI 1640 medium to prepare single-cell suspensions. Red blood cells were removed using red blood cell lysis buffer (Sigma Aldrich) according to the manufacturer’s instructions. For surface marker analysis, leukocytes were treated with FcγR blocker CD16/32 (2.4G2, BD) and then incubated with fluorochrome-labeled antibodies or viability dye for 30 min. For analysis of intracellular markers, cells were stimulated with either ZIKV peptide (Env294–302, 5 ug/ml) or PMA (50 ng/ml)/ionomycin (750 ng/ml), in both cases with the addition of brefeldin, for 5 h, followed by the fixation and permeabilization using IC buffer (Thermo Fisher Scientific). For analysis of CD8^+^ IFN-γ^+^ or CD8^+^ TNF^+^ T cells, cells were stimulated with ZIKV peptide and for analysis of CD4^+^ IFN-γ^+^ or CD4^+^ IL-17^+^ T cells, cells were stimulated with PMA/Ionomycin. The fluorochrome-labeled antibodies used were purchased from Thermo Fisher Scientific, Tonbo Biosciences or Biolegend as below: Live Dead Fixable Dye (eFluor 506), PE-Cy7 anti-CD3 (clone 145-2C11), violetFluor 450 anti-CD8 (Clone 53-6.7), PerCP-Cy5.5 anti-CD4 (clone RM4-5), FITC anti-CD4 (Clone GK1.5), BV711 anti-CD44 (IM7), APC anti-CD62L (Clone MEL-14), FITC anti-CXCR3 (Clone CXCR3-173), PerCP-Cy5.5 anti-CD11b (Clone M1/70), BV421 anti-F4/80 (Clone T45-2342), APC anti-Ly6G (Clone 1A8), PE-Cy7 anti-CD80 (16-10A1), FITC anti-CD11c (Clone N418), APC anti-IFN-γ (Clone XMG1.2), PE anti-PD-1 (clone J43), PerCP-eflour 710 anti-TNF (Clone MP6-XT22), BV711 anti-IL-4 (Clone 11b11), and PE anti-IL-17 (Clone ebio17b7). To inactivate the virus, cells were fixed in paraformaldehyde 2% for 16 h at 4°C. Cell samples were acquired on BD LSR Fortessa and data were analyzed (Figure S1) using FlowJo 10 (BD).

### 4.5 Histology

Samples of tissue were collected and placed into a solution of 10% neutral buffered formalin (Thermo Fisher Scientific) for 24 h. Following this incubation, the formalin solution was replaced with 95% ethanol and were either stained with hematoxylin-eosin (H&E) or left unstained and used for immunofluorescence analyses. A blinded board-certified pathologist read the H&E slides.

### 4.6 Immunofluorescence assay

To conduct immunofluorescence assays, a modified protocol from a previous study was used ^42^. Sections of mouse testes on slides were dewaxed by heating at 65 °C overnight, followed by incubations in xylene and graded ethanol series (Millipore, Sigma). Antigen retrieval was performed by incubation in citrate buffer pH 6 (Abcam) at 90°C for 30 min. Sections were then washed in deionized water and blocked with a solution of 5% normal goat serum (NGS) in PBS for 15 min at 37 °C and then stained overnight at 4 °C with 4G2 antibody (Abcam) at 1:200. Washed in PBS and incubated with 0.2% Sudan black B diluted in 70% ethanol for 12 min at 37°C. The sections were then incubated for 45 min at 37°C with a secondary antibody at a dilution of 1:2000, AlexaFluor 594 anti-human IgG (Thermo Fisher). The slides were mounted with ProLong Gold Antifade Mountant with DAPI (Thermo Fisher). Images were captured using an inverted fluorescence microscope (Olympus-IX73).

### 4.7 Quantification and statistical analyses

Mouse groups weight data were compared at each time point using two-way ANOVA with Tukey’s multiple comparison correction. For comparing ZIKV positive/negative immunofluorescence of the testes, Fisher’s exact test was used. For all other statistical tests, either one-way ANOVA with Sidak’s multiple comparison testing was used for comparison between multiple groups or two-tailed student’s t-test was used when only two groups were being compared. Viral titers were log_10_ transformed before statistical tests were applied. Left and right testicular weights and percent of spermatogenesis were averaged before statistical analysis. In the experiment presented in Figure S4, only left testis data was available because the right testis was used for titration data, and cutting the testes was found to reduce the quality of the histology data. Data from flow cytometry, BioPlex, testicular weight, and percent spermatogenesis were compared using one-way ANOVA with Sidak’s multiple comparison testing. For all analyses, a p-value lower than 0.05 was considered significant.

## DATA AVAILABILITY

All data is available upon reasonable request.

## AUTHOR CONTRIBUTIONS

Conceptualization: R.K.C., J.S., Y.L., and S.L.R. Formal analysis: R.K.C., Y.L., J.S. and S.L.R. Investigation: R.K.C., Y.L., S.R.A., J.L., V.N.C., E.E.H.S., and S.L.R. Resources: J.S. and S.L.R. Data curation: R.K.C., Y.L., and S.R. Writing-original draft preparation: R.K.C., J.L., S.L.R. Writing-review and editing: R.K.C., Y.L., S.R.A., J.L., V.N.C., E.E.H.S., J.S., and S.L.R. Project administration: S.L.R. All authors contributed to the article and approved the submitted version.

## ACKNOWLEDGMENTS

Viral isolates used in this study were provided by the World Reference Center for Emerging Viruses and Arboviruses (WRCEVA) at the University of Texas Medical Branch. Graphs and figures were created with Graphpad prism version 10 and BioRender.com, respectively.

---

**Figure S1.**
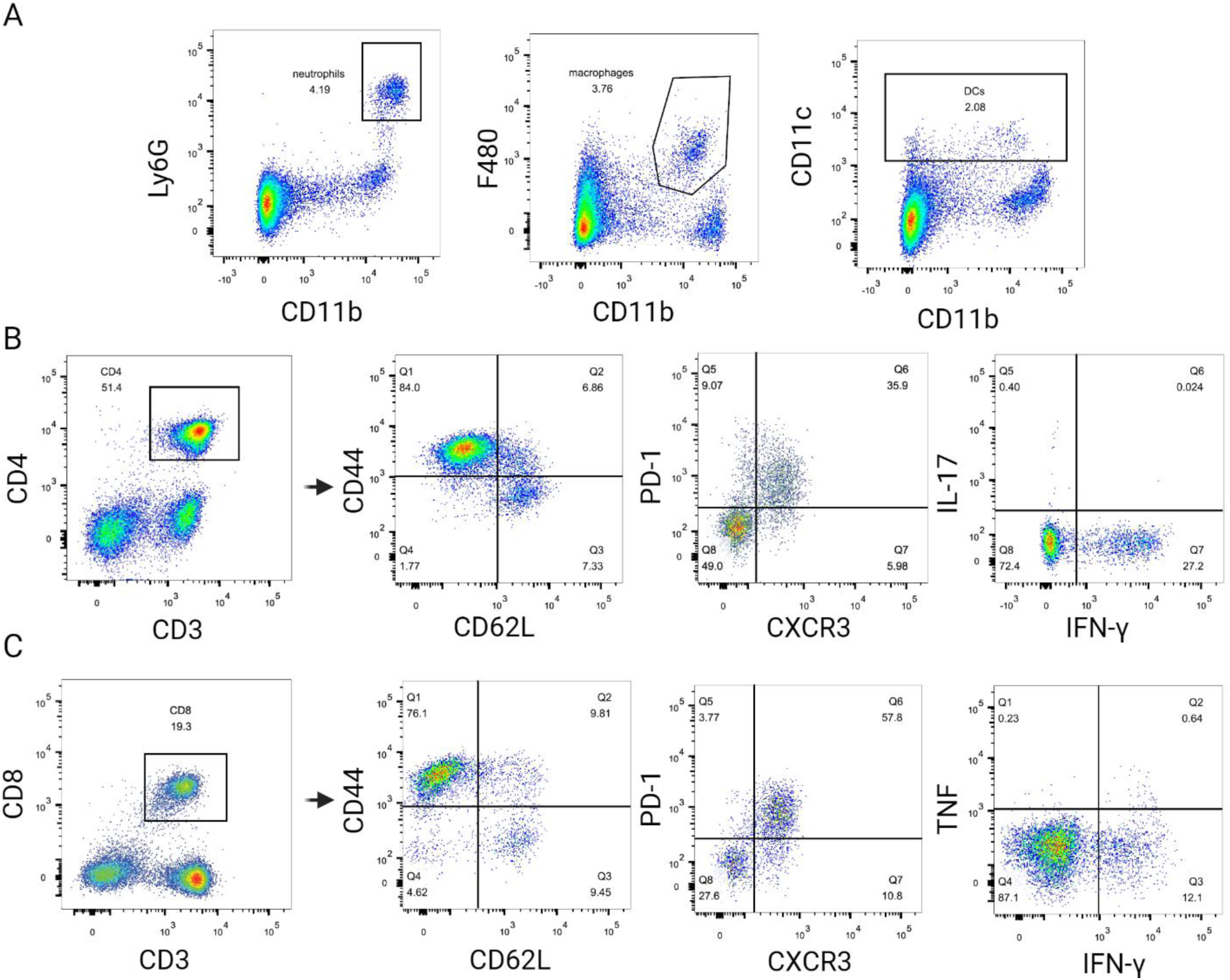
Flow cytometry gating strategy. Cells were first gated by side scatter and forward scatter, singlets were selected, live/dead-negative cells were selected. A. Neutrophils (Ly6G^+^ CD11b^+^), macrophages (F480^+^ CD11b^+^) and dendritic cells (CD11c^+^) gating strategy. B. CD4^+^ T cells gating strategy. These cells were then selected and gated for effector status (CD44^hi^ CD62L^lo^), PD-1, CXCR3, IL-17 and IFN-gamma. C. CD8^+^ T cells gating strategy. These cells were then selected and gated for effector status (CD44^hi^ CD62L^lo^), PD-1, CXCR3, TNF and IFN-γ.

**Figure S2:**
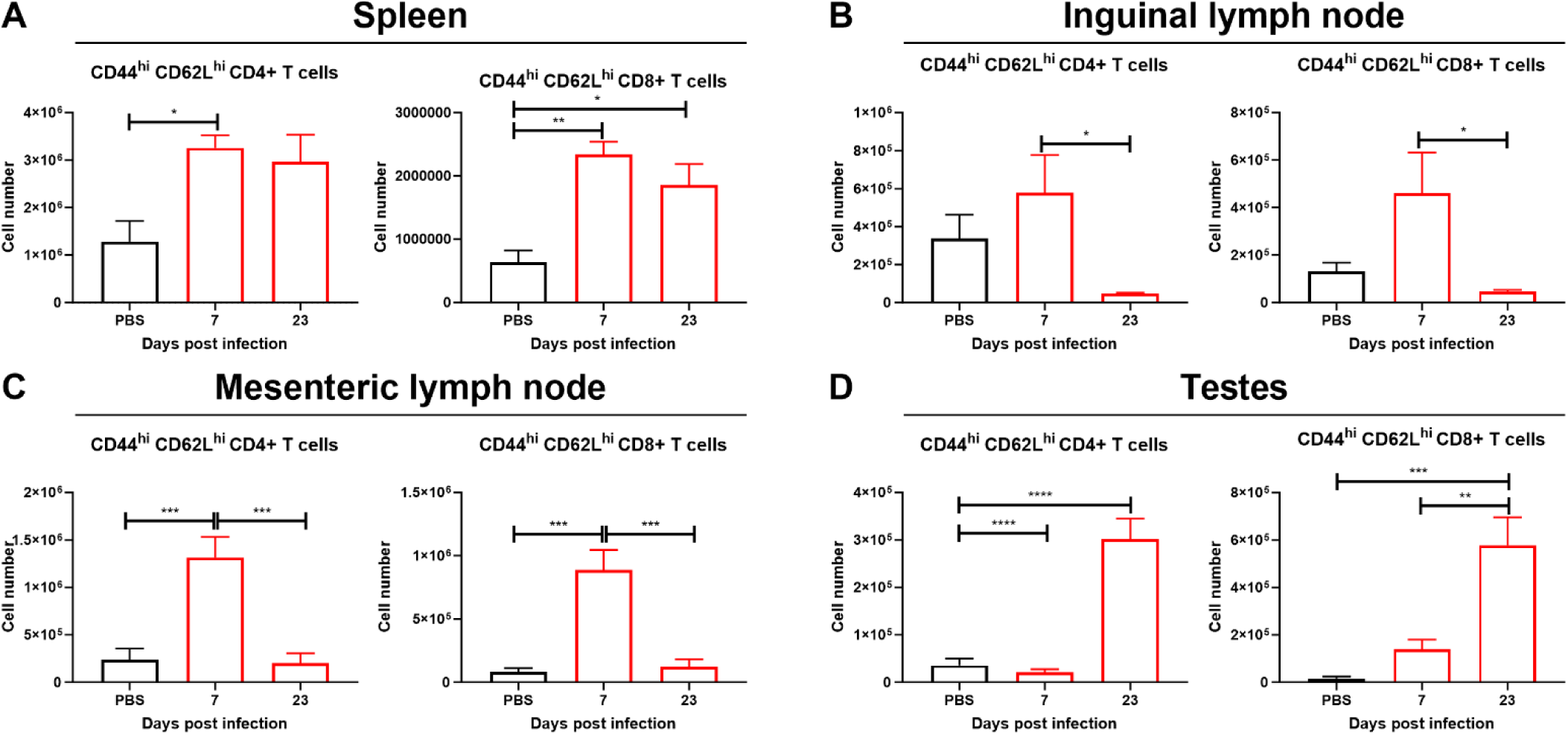
CD44^hi^ CD62L^hi^ T cells are present in testes at 23 dpi. Possible memory CD8^+^ and CD4^+^ T cells detected in A. the spleen, B. the inguinal lymph nodes, C. the mesenteric lymph nodes, D. the testes. Red bars show ZIKV-infected mice, black bars denote uninfected baseline controls. Statistical significance was assessed with one-way ANOVA Sidak’s multiple comparison correction. Bars show mean and error bars represent standard error of the mean. *, p < 0.05; **, p < 0.01; ***, p < 0.001; ****, p < 0.0001.

**Figure S3.**
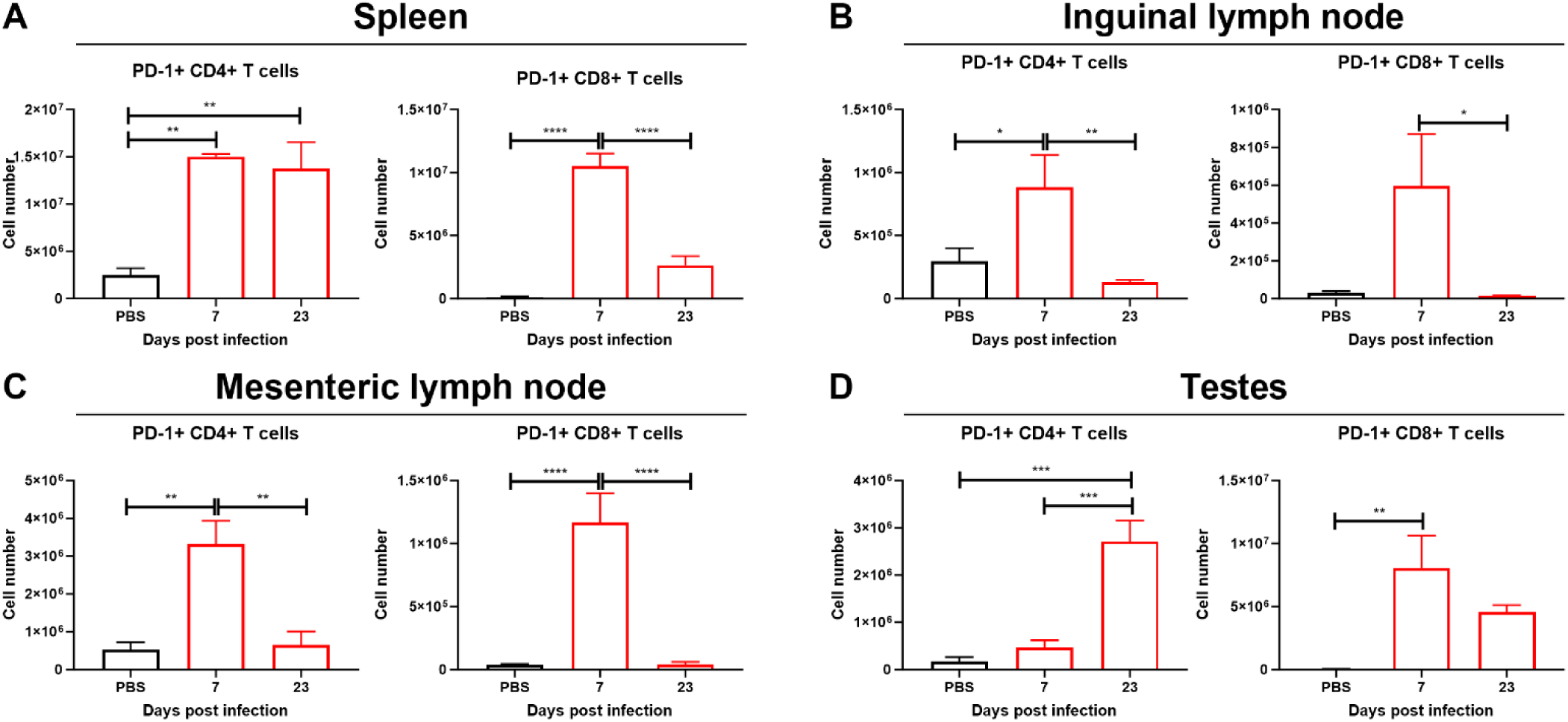
Analysis of PD-1^+^ T cells. Cell number of PD-1+ T cells in A. the spleen, B. the inguinal lymph nodes, C. the mesenteric lymph nodes, D. the testes. Red bars show ZIKV-infected mice, black bars denote uninfected baseline controls. Statistical significance was assessed with one-way ANOVA Sidak’s multiple comparison correction. Bars show mean and error bars represent standard error of the mean. *, p < 0.05; **, p < 0.01; ***, p < 0.001; ****, p < 0.0001.

**Figure S4:**
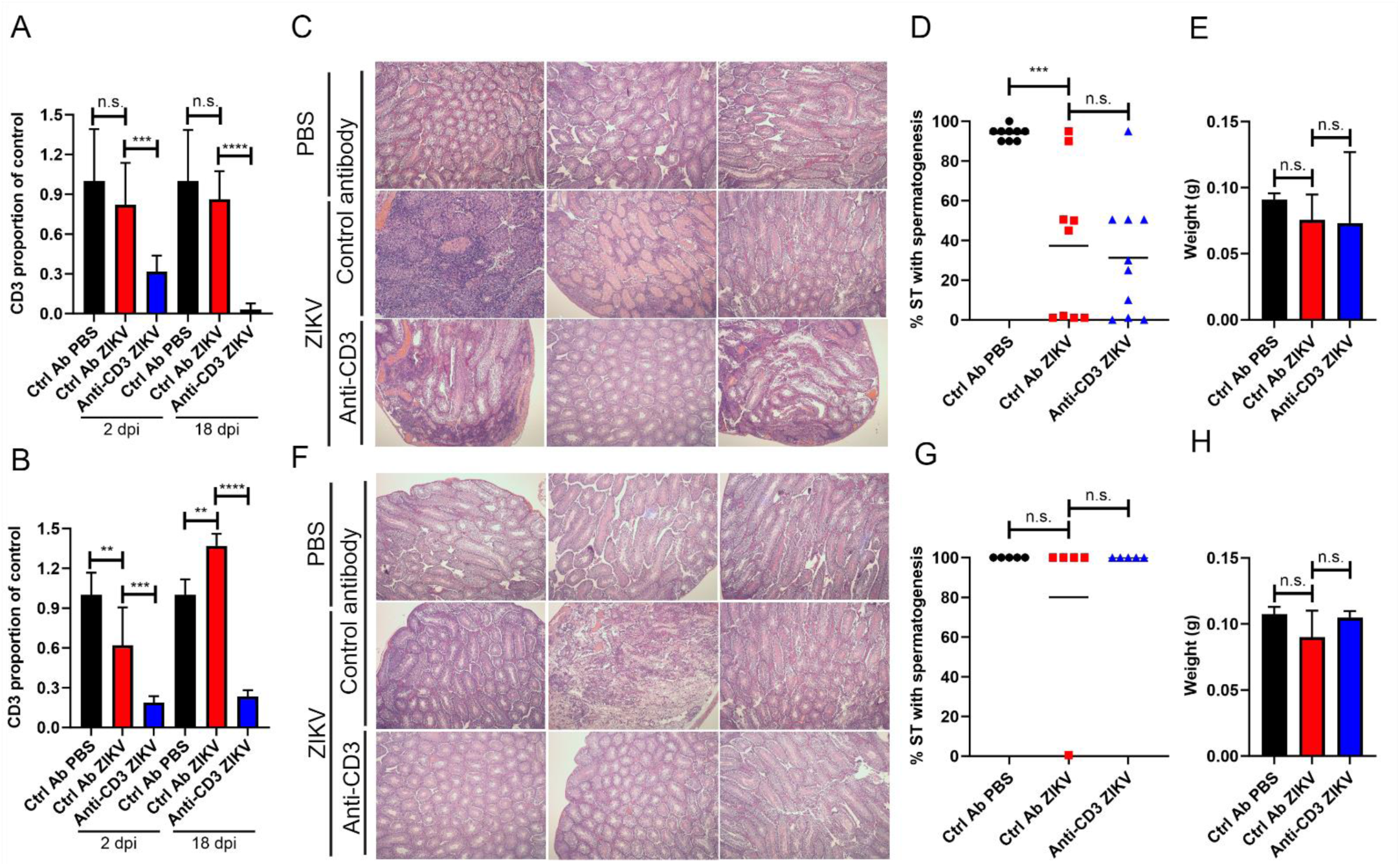
Depletion of CD3 did not significantly impact testicular damage of A129 mice. A. Histology of A129 mice testes. B. Percentage of CD3 events in blood samples of A129 mice excluding red blood cells at 2 or 18 dpi. C. Percentage of seminiferous tubules (ST) with spermatogenesis in each A129 mouse testicular sample. D-F show histology, CD3 levels, and percentage of ST for the C57BL/6J mice. Statistical significance was assessed with one-way ANOVA Sidak’s multiple comparison correction. Bars show mean and error bars represent standard deviation. Lines represent the mean. **, p < 0.01; ***, p < 0.001; ****, p < 0.0001; n.s., no significant difference.

